# Multiple repeat regions within mouse DUX recruit chromatin regulators to facilitate an embryonic gene expression program

**DOI:** 10.1101/2023.03.29.534786

**Authors:** Christina M. Smith, Edward J. Grow, Sean C. Shadle, Bradley R. Cairns

**Affiliations:** Howard Hughes Medical Institute, Department of Oncological Sciences and Huntsman Cancer Institute, University of Utah School of Medicine, Salt Lake City, UT, USA; Green Center for Reproductive Biological Sciences, Department of Obstetrics and Gynecology, University of Texas Southwestern Medical Center, Dallas, TX, USA

## Abstract

The embryonic transcription factor DUX regulates chromatin opening and gene expression in totipotent cleavage-stage mouse embryos, and its expression in embryonic stem cells promotes their conversion to 2-cell embryo-like cells (2CLCs) with extraembryonic potential. However, little is known regarding which domains within mouse DUX interact with particular chromatin and transcription regulators. Here, we reveal that the C-terminus of mouse DUX contains five uncharacterized ∼100 amino acid (aa) repeats followed by an acidic 14 amino acid tail. Unexpectedly, structure-function approaches classify two repeats as ‘active’ and three as ‘inactive’ in cleavage/2CLC transcription program enhancement, with differences narrowed to a key 6 amino acid section. Our proximity dependent biotin ligation (BioID) approach identified factors selectively associated with active DUX repeat derivatives (including the 14aa ‘tail’), including transcription and chromatin factors such as SWI/SNF (BAF) complex, as well as nucleolar factors that have been previously implicated in regulating the Dux locus. Finally, our mechanistic studies reveal cooperativity between DUX active repeats and the acidic tail in cofactor recruitment, DUX target opening, and transcription. Taken together, we provide several new insights into DUX structure-function, and mechanisms of chromatin and gene regulation.

## Introduction

Mouse pre-implantation development is a dynamic and complex process, which begins with a single-cell zygote and results in an implanted embryo bearing cells with different lineage specifications and fates. Within this period, numerous molecular processes must be properly completed, which are driven (in part) by gene expression changes and chromatin modifications (1,2). A hallmark of pre-implantation development is major embryonic genome activation (EGA), which occurs at the 2-cell stage in the mouse embryo(3). This critical process includes decreases in maternally-deposited mRNAs and proteins, combined with the onset of both embryo-derived transcription and translation – effectively turning developmental control over to the embryo from maternal resources in the egg (4). During EGA, chromatin undergoes considerable remodeling to enable enhancer and promoter accessibility, allowing for the onset of major embryonic transcription, and the marking of promoters and enhancers for future regulation (5–7).

Mouse embryonic stem cells (mESCs) are derived from the inner cell mass (ICM) of the blastocyst, and have enabled studies focused on the characterization of the blastocyst and downstream embryonic differentiation(8). In addition, the mESC system has been extended to examine properties of earlier cleavage-stage totipotent-related cells. Specifically, a small (<1%) population of spontaneously fluctuating mESCs has been identified, which closely resemble cells within the 2-cell embryo, termed 2-cell-like cells (2CLC) (9–12). Our lab and others showed that the expression of Dux, which encodes the double homeodomain transcription factor DUX, is sufficient to confer this 2C-like state to mESCs (12–14). However, we currently have an incomplete understanding regarding how DUX establishes the 2C transcriptome. DUX4, the human Dux ortholog is similarly expressed during EGA, which occurs at the 4-cell stage in humans. Likewise, DUX4 expression in human ESCs converts these cells to transcription programs resembling the 4-8 cell human cleavage-stage embryo (13,15–17). DUX4 has additionally been identified as the causal gene in facioscapulohumeral dystrophy (FSHD), leading to the protein being intensely studied to identify its pathological traits (18–20).

Mouse DUX is known to be targeted to specific sites in the genome (including genes and MERVL LTRs) by its two N-terminal homeodomains, whereas its C-terminal region drives transcriptional activation(12,21,22). However, little is known about the nature of this C-terminal region, which is much more divergent among DUX orthologs than are the homeodomains (23,24). One possible contributor to transcriptional activation involves the interaction of the C-terminus of DUX4 with the related histone acetyltransferases p300 and CBP (22,25). However, these or other interactions have not been identified with mouse DUX. Regarding chromatin remodeling during EGA, the abundant ATP-dependent chromatin remodeler SWI/SNF (BAF) complex plays a central role(26,27). Specifically, SMARCA4, the catalytic ATPase within SWI/SNF complex, is required for important for EGA in the 2-cell mouse embryo (28). SMARCC1 is a core structural scaffold for SWI/SNF complexes, and its absence results in loss of complex activity *in vivo* (29). However, possible physical links between mouse DUX and the chromatin remodeler SWI/SNF complex during EGA or during the mESC to 2C-like transition is currently unknown.

DUX activates about 25% of EGA genes and DUX expression is sufficient to revert mESCs to an earlier totipotent stage(12). Consistent with DUX impacting early development, *Dux-/-* crosses yield smaller litter sizes (21). However, as viable Dux -/-pups are generated, factor(s) redundant with DUX appear to contribute to DUX target activation and chromatin reprogramming in 2-cell embryos (21,30). Recently, oocyte-specific homeobox 4 (OBOX4), a factor highly expressed during EGA and in 2C-like mESCs, has been identified as partly redundant with DUX in 2-cell embryos(31). Taken together, DUX and OBOX4 appear to cooperate to ensure proper EGA in the early mouse embryo.

In this study, we sought to better characterize DUX domains, identify proteins interacting with DUX domains, and understand how DUX domains contribute to DUX target gene activation, DUX promoter recruitment, and chromatin accessibility at binding sites. Specifically, we reveal a unique C-terminal repeat structure in mouse DUX, and characterize two functionally distinct repeat types (termed ‘active’ and ‘inactive’) and cooperativity of the active repeats with the C-terminal acidic tail. Additionally, we identify DUX domains that interact with the chromatin remodeler subunit SMARCC1, which alongside the ability of DUX to confer histone acetylation, reveals new mechanistic insight into how the C-terminus of DUX confers transcriptional activation.

## Results

### The mouse DUX C-terminus has five conserved repeats followed by a 14 amino acid ‘tail’

Curiously, mouse DUX protein (673 amino acids (aa)) is considerably longer than DUX orthologs such as human DUX4 (424 aa). Through manual inspection and alignments of mouse DUX with other orthologs, we identified five protein repeats within DUX of ∼100 aa each, followed by a single 14aa highly-acidic C-terminal ‘tail’ (Figure 1a). However, human DUX4 contains only a single copy of this ‘repeat’ domain, which is likewise followed by a 14aa tail that closely resembles the acidic tail in mouse DUX (Figure 1b). Prior work has revealed that the acidic DUX4 C-terminal tail helps activate transcription, in part by binding to CBP and p300 (22). To determine whether other placental mammals (e.g. rat, bovine, and human) contain a C-terminal repeat structure similar to mouse DUX, we conducted pairwise alignments using the algorithm BLAST, which revealed repeats only in the mouse lineage (Figure 1c, Supplementary Figure 1a,b,c). Interestingly, although all five repeats generate highly significant pairwise similarity scores, two pairs with the highest similarity scores involve the C3-C5 pair and the C2-C4 pair – results supported by our earlier phylogenetic analyses (Figure 1c). Furthermore, the human DUX4 C-terminal region displays significant similarity to both the 5^th^ and 2^nd^ repeat of mouse DUX (E-Values 0.022 and 0.03, respectively). The bovine DUXC C-terminal region was most similar to the 5^th^ repeat whereas the rat C-terminus was most similar to the 1^st^ repeat of mouse DUX (Supplementary Figure 1d). However, as the comparisons DUX C-terminal ‘single repeat’ orthologs to the mouse repeats yield different mouse repeats with highest similarity, these results do not point clearly to a particular repeat representing the parental/original repeat in mouse DUX.

**Figure 1:**
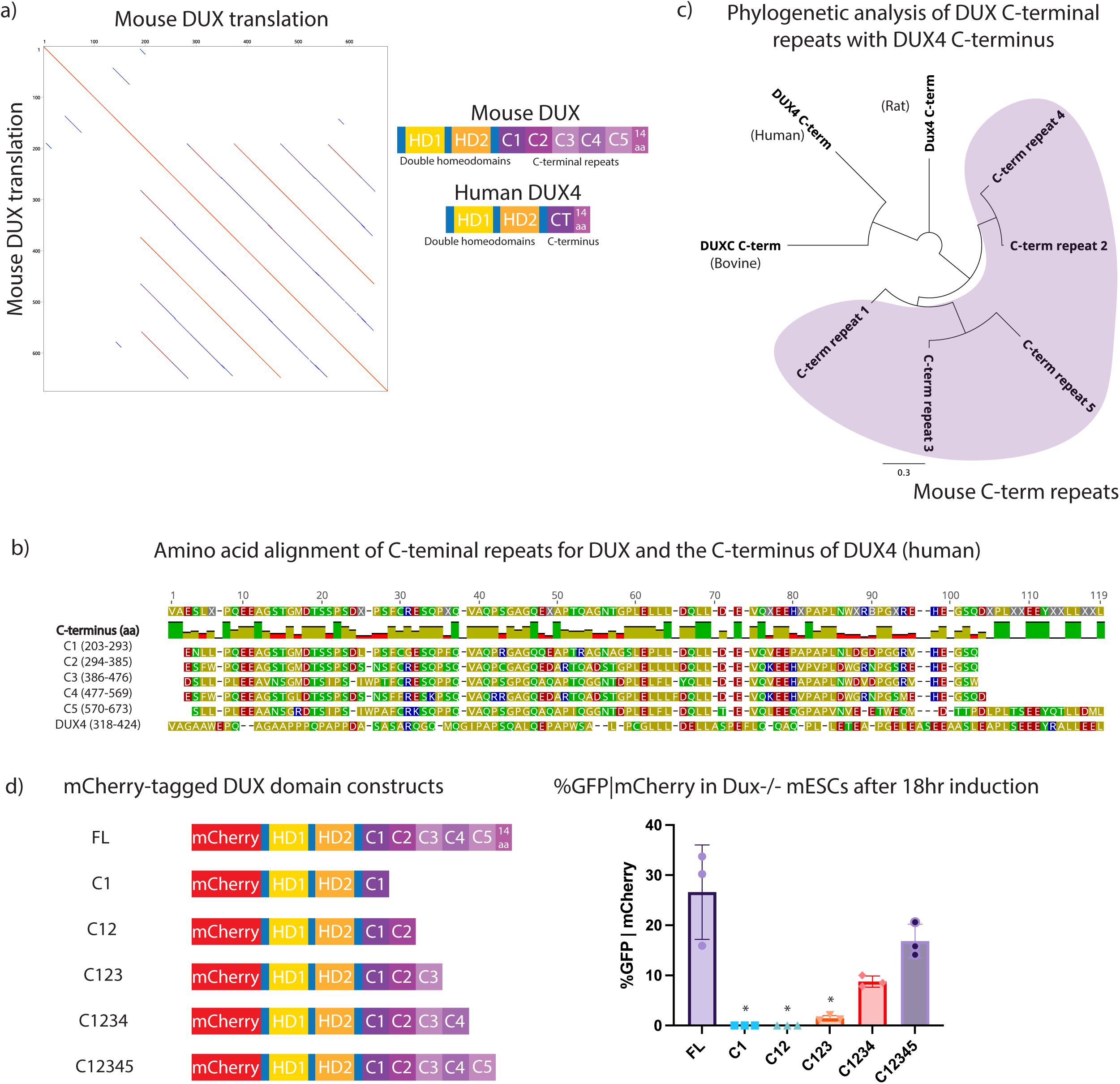
Identification of five repeats within the mouse DUX C-terminal region, and truncation analysis of transcriptional activity. a. (Left) Mouse DUX aligned to itself, revealing five repeats. (Right) Diagram of mouse DUX and human DUX4. b. Amino acid alignment of mouse DUX C-terminal repeats and comparison to human DUX4. A 14 amino acid tail follows both the terminal mouse C5 repeat and the sole DUX4 repeat. c. Phylogenetic analysis showing amino acid alignments of mouse DUX C-terminal repeats vs. the C-terminus of bovine DUXC, the C-terminus of rat DUX4, and the C-terminus of human DUX4. Line length unit is substitutions per base. Purple shading denotes mouse DUX C-terminal repeats. d. (Left) Schematic of constructs used for flow cytometry: mCherry-tagged full-length DUX (FL), homeodomains with C1, C12, C123, C1234, C12345. (Right) Flow cytometry data for MERVL::GFP reporter given mCherry expression in *Dux-/-* mESCs following 18hr expression of indicated constructs. *p-value < 0.05, student’s t-test. (n=3 biological replicates)

Next, we assessed the contribution of mouse DUX repeats to transcriptional activity by examining *Dux-/-* mESCs containing a MERVL::GFP reporter. The long-terminal repeat (LTR) of MERVL is bound by DUX, and MERVL:GFP is commonly used to assess DUX activation and conversion of mESCs to a 2CLC state (9,12). First, *Dux-/-* mESCs were transiently transfected with mCherry-tagged DUX repeat domain constructs encoding zero to five C-terminal repeats, without the 14aa tail (Figure 1d). Cells were induced with doxycycline for 18 hours, gated on mCherry positivity, and MERVL::GFP expression was quantified by flow cytometry (Supplementary Figure 1e). Expression of each derivative was confirmed by Western analysis (Supplementary Figure 1f). The flow cytometry results reveal that the first two repeats have minimal activity, but with the addition of the third repeat, low MERVL::GFP expression is observed. Activity is further improved by the addition of the fourth and fifth repeat. The results also support a potential role for the 14aa ‘tail’ in activation, as its omission modestly lowered MERVL::GFP expression (Figure 1d). Thus, our initial deletion series revealed activity with repeats 3-5, but not with repeats 1-2 alone. However, this experiment did not determine whether the repeats act in a simple additive manner, or if individual repeats confer distinct transcriptional activity in a singular manner, or how they cooperate with the acidic tail.

### DUX C-terminal repeats C3 and C5 contribute to transcriptional activation when combined with the 14aa tail

We next explored the contributions of individual repeats, along with their cooperation with the C-terminal acidic ‘tail’. Here, we generated N-terminal mCherry-tagged DUX derivatives with varying numbers of repeats either with or without the 14aa tail, as well as with or without single or both homeodomains (Supplementary Figure 2a). All constructs contain a GGGGS_2_ linker present in constructs with a single internal deletion and GGGGS_2_ and GAGAS_2_ with constructs with two internal deletions. First, as expected, derivatives lacking either homeodomain failed to activate the MERVL::GFP reporter (Supplementary Figure 2a). Second, combinations involving C1, C2 or C4 with the acidic ‘tail’ lack activation of the MERL::GFP reporter, whereas combinations of C3 or C5 with the ‘tail’ display potent reporter activation (Figure 2a). The latter combinations require the 14aa tail as constructs lacking the tail, but containing the C3 or C5 repeat alone, did not activate the reporter. Notably, a construct containing C1 and C2 with the 14aa tail was inert, whereas C1, C2 and C3 with the 14aa tail was highly active (Supplementary Figure 2a,b). Together, these results suggest that strong DUX target gene activation requires a combination of two C-terminal domains: a transcriptionally ‘active’ repeat (C3 or C5) and the 14aa tail (Figure 2b).

**Figure 2:**
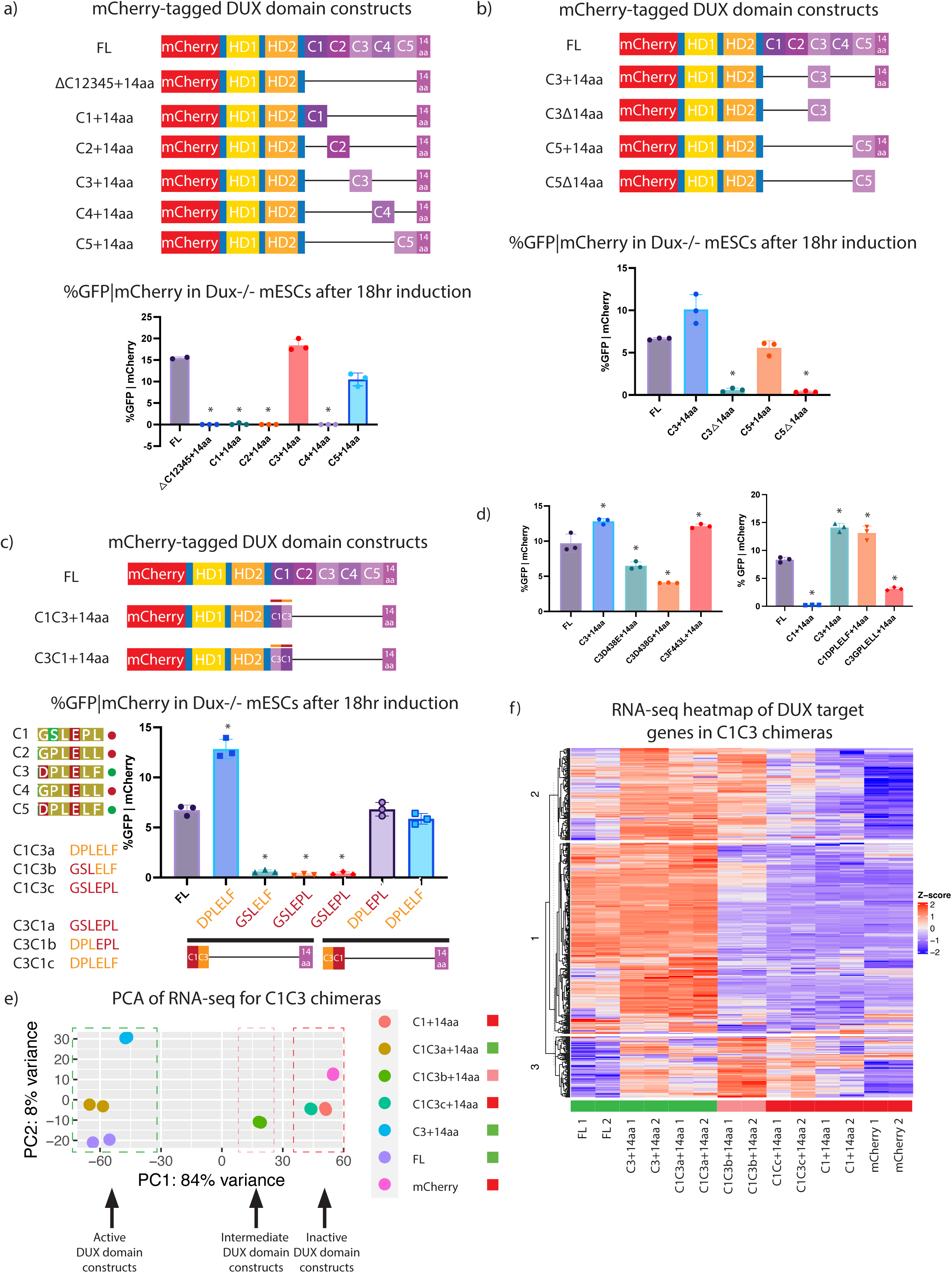
Structure function-analysis of DUX by 2C-like cell conversion. a. (Top) Schematic of constructs used for flow cytometry: mCherry-tagged FL, ΔC12345+14aa, C1+14aa, C2+14aa, C3+14aa, C4+14aa, and C5+14aa. (Bottom) Flow cytometry for MERVL::GFP reporter given mCherry expression in *Dux-/-* mESCs with 18hr overexpression of indicated constructs. *p-value < 0.05, student’s t-test. (n=3 biological replicates). Constructs with a single black line have the linker GGGGS_2_ and constructs with two black lines have linkers with GGGGS_2_ and GAGAS_2_, respectively. b. (Top) Schematic of constructs used for flow cytometry: mCherry-tagged FL, C3+14aa, C3Δ14aa, C5+14aa, C5Δ14aa. (Bottom) Flow cytometry for MERVL::GFP reporter given mCherry expression in *Dux-/-* mESCs following 18hr expression of indicated constructs. *p-value < 0.05, student’s t-test. (n=3 biological replicates) c. (Top) DUX domain chimera construct design for C1-C3 fusions with (top) schematic of constructs and (bottom) specific chimera cut-offs for C1-C3 a,b,c (red to orange and C3-C1 a,b,c (orange to red) constructs. (Bottom) Schematic of DUX domain chimera constructs used for flow cytometry: mCherry-tagged full-length DUX (FL), C1C3a, C1C3b, C1C3c, C3C1a, C3C1b, C3C1c. Symbols adjacent to alignments: red circle denotes transcriptionally inactive repeats and the green circle denotes transcriptionally active repeats. (Right) Flow cytometry for MERVL::GFP reporter given mCherry expression in *Dux-/-* mESCs following 18hr expression of indicated constructs. *p-value < 0.05, student’s t-test. (n=3 biological replicates) d. Flow cytometry for MERVL::GFP reporter given mCherry expression in *Dux-/-* mESCs following 18hr expression of point mutation constructs: (Left) FL-DUX, C3+14aa, C3 D438E+14aa, C3 D438G+14aa, and C3F443L+14aa. (Right) FL-DUX, C1+14aa, C3+14aa, C1DPLELF+14aa (substation of C3 amino acids into C1+14aa), and C3GPLELL+14aa (substitution of C2 and C4 6aa sequence into C3+14aa). *p-value < 0.05, student’s t-test. (n=3 biological replicates) e. Principal component analysis (PCA) of RNA-seq analysis for 18hr overexpression of DUX domain chimera constructs, full-length DUX, and mCherry alone (n=2). Colored squares beside sample legends denote the inability to activate MERVL::GFP reporter (red) or ability to activate MERVL::GFP reporter (green). f. Heat map of RNA-seq at DUX target genes (n=456) from 18hr expression of DUX domain chimera constructs, full-length DUX, and mCherry alone (n=2). Colored rectangles above samples denote inability to activate MERVL::GFP reporter (red) or ability to activate MERVL::GFP reporter (green).

To determine the amino acids selectively involved in DUX target activation within the active C3 and C5 repeats, chimeric constructs between C1 (a transcriptionally ‘inactive’ repeat) and C3 (a transcriptionally ‘active’ repeat) were made, all containing the 14aa tail. The cutoff/transition point between chimera regions occurred in 3 aa increments along the 2 repeats (Figure 2c, Supplementary Figure 2c). This narrowed down the functional difference to a 6 aa region which is conserved between repeats C3 and C5, but not conserved in C1, C2, and C4. Importantly, replacement solely of this 6 aa sequence from the active C3 repeat into inactive C1 repeat (termed C1DPLELF+14aa) conferred robust activation of the MERVL::GFP reporter (Figure 2d).

In order to expand these analyses to global transcription and all DUX targets (beyond the MERVL::GFP reporter), we performed RNA-seq on full-length (FL) DUX and selected chimera constructs. This approach was designed to test whether the ‘active’ constructs might differ in their gene targets (Figure 2e and Supplementary Figure 2d). First, the PCA plot for these samples separated samples largely into two groups – active versus inactive – with one sample in an intermediate position (Figure 2e). Interestingly, the DUX constructs which activate the MERVL::GFP reporter (FL, C1C3a, and C3+14a) all showed strong activation of DUX target genes, whereas the ‘inactive’ constructs that do not activate the MERVL::GFP reporter (C1+14aa, C1C3c, and mCherry alone) did not activate DUX target genes (Figure 2f). Notably, the intermediate position was occupied by the C1C3b derivative, reinforcing the notion that the 6aa region is important for regulating target activation. K-means clustering of the RNA-seq for the chimera constructs at DUX target generated three clusters of genes: the first two of which largely contained genes activated by FL DUX, whereas the cluster #3 contains genes that have increased expression in DUX domain derivatives compared to FL DUX. The ‘intermediate’ derivative C1C3b, which is a hybrid of an ‘active’ and an ‘inactive’ repeat, was capable of activating the majority of DUX targets in cluster #2, but failed to activate those in cluster #1, whereas it activated (or failed to attenuate) genes in cluster #3. Genes activated by C1C3b in cluster #3 generate GO terms such as ‘upregulated include synapse assembly’, ‘sex differentiation’, ‘reproductive structure development’, and ‘neuron differentiation’. Here, cluster 3 shows examples of known DUX target genes that have a higher relative expression in C1C3b compared to full-length DUX include Obox1, *Olfr1277, Obox2*, and *Zfp296*. These combined observations, especially those involving derivatives that cause the precocious activation of genes in cluster #3, raise the interesting possibility that the ‘inactive’ repeats may serve to attenuate the active repeats at particular targets, such as *Obox* genes. Finally, when considering amino acid similarity between the five repeats, we note that C3 and C5 are most similar to one another, compared to C1, C2, and C4 (Supplementary Figure 2e).

### Chromatin alterations at DUX binding sites are associated with a transcriptionally active DUX repeat

To investigate the chromatin landscape effects conferred by Dux derivative expression, we performed CUT&Tag in *Dux-/-*mESCs on selected Dux domain constructs: FL, C12345△14aa, C3+14aa, C3△14aa, △C12345+14aa, and mCherry alone. Because all of the constructs had the same N-terminal mCherry tag, an mCherry antibody was used throughout. We isolated and analyzed clonal stable cell lines which expressed species of the expected protein size and demonstrated comparable expression to one another (Figure 3a). We note that mCherry-DUX protein is expected to run at roughly 100kDa, but runs anomalously on polyacrylamide gels – at 150kDa. This is likely due to a property of the C-terminal repeats as a DUX derivative consisting of the homeodomains alone runs at the expected molecular weight. To benchmark DUX CUT&Tag against previous chromatin occupancy data, we first generated a CUT&Tag dataset for mCherry-tagged full-length DUX protein, and found that it overlapped highly with previously published HA-tagged DUX ChIP-seq results (Supplementary Figure 3a) (12). Extension of CUT&Tag to the DUX derivatives revealed their moderate-to-high occupancy at DUX sites, except for C3△14aa, which displayed reduced occupancy at DUX binding sites (Figure 3b, 3c). Next, we determined the ability of our DUX domain constructs to alter H3K9ac, as H3K9ac globally increases in 2C-like cells (32). Notably, H3K9ac occurs at DUX binding sites correlating with the transcriptional activity of each Dux construct (Figure 3d,e, Supplementary Figure 3b). Taken together, our data demonstrate that the 14aa acidic tail is not strictly required for the occupancy of DUX targets. Furthermore, our data suggest that the 14aa tail is important for conferring H3K9ac at bound sites and might also be involved in retention of DUX at target sites since removal of the 14aa tail (in the context of a single active repeat) diminished target occupancy.

**Figure 3:**
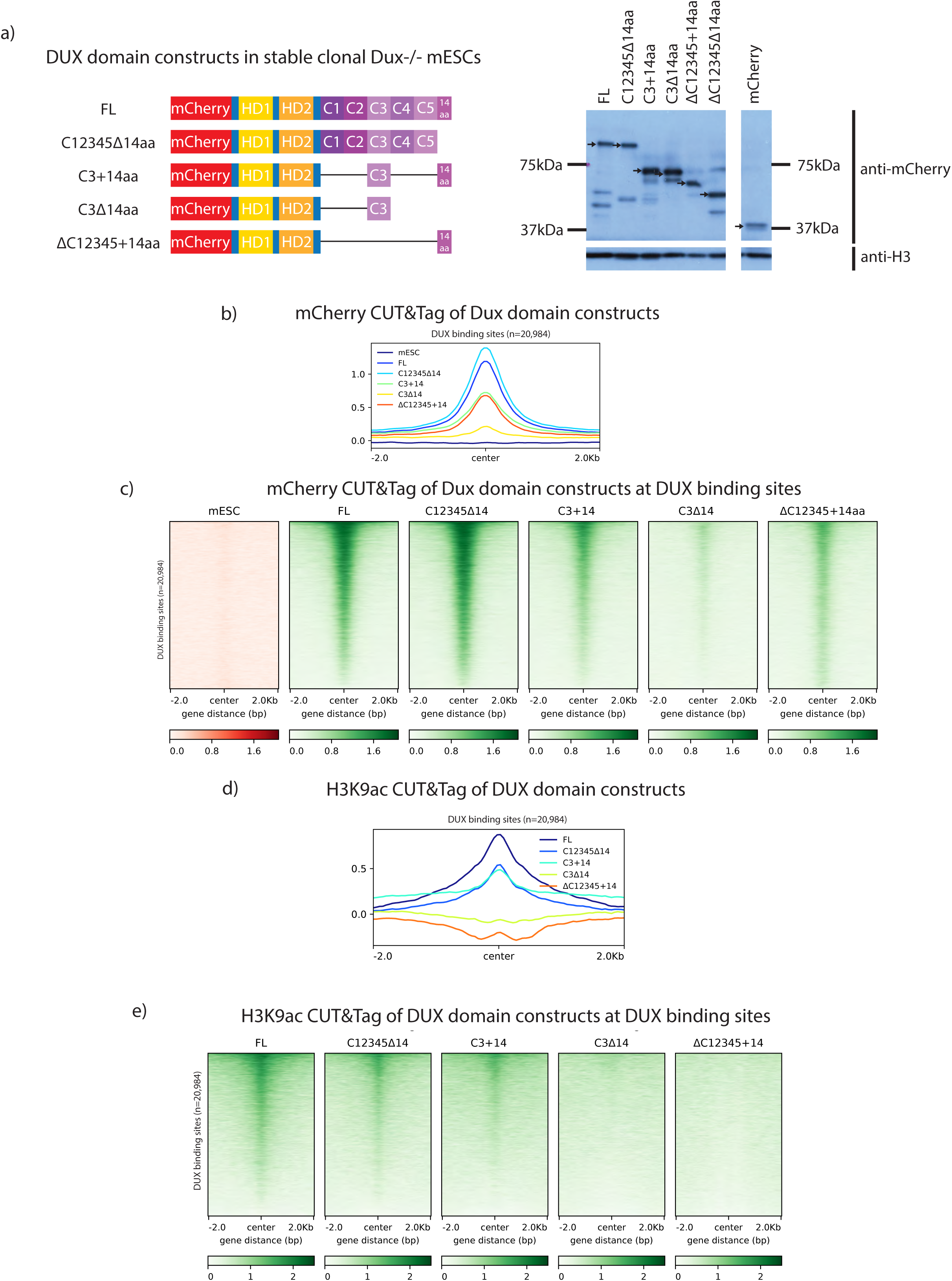
Chromatin modifications at DUX binding sites requires an active DUX repeat. a. (Left) Schematic of mCherry-tagged DUX domain constructs in clonal *Dux-/-* mESCs used for CUT&TAG experiments. (Right) anti-mCherry Western blot for indicated constructs after 18hr expression. Black arrows indicate construct of interest. Predicted protein sizes are 100 kDa (FL), 95 kDa (C12345Δ14aa), 65 kDa (C3+14aa), 59 kDa (C3Δ14aa), 48 kDa (ΔC12345+14aa), and 25 kDa (mCherry). b. mCherry CUT&TAG class average map centered at DUX binding sites after 12-hour overexpression of Dux constructs illustrated in Figure 2a (n=2 biological replicates). c. mCherry CUT&TAG heatmap centered at DUX binding sites after 12-hour expression of Dux constructs illustrated in Figure 2a (n=2 biological replicates). d. H3K9ac CUT&TAG class average map centered at DUX binding sites after 12-hour expression of Dux constructs illustrated in Figure 2a (n=2 biological replicates). e. H3K9ac CUT&TAG heatmap centered at DUX binding sites after 12-hour expression of Dux constructs illustrated in Figure 2a (n=2 biological replicates).

### Proximity-labeling with full-length DUX reveals interactions with proteins involved in chromatin de-repression

Next, we identified proteins that interact with DUX derivatives using the BioID approach, which identifies proteins that interact directly, indirectly, or transiently with a protein of interest (33). Here, we utilized BioID to define protein interactions with DUX in the 2C-like cellular environment. To accomplish this, we fused BirA*, a promiscuous biotinylating enzyme, to the N-terminus of full-length DUX. When biotin is added to the growth media, BirA* will biotinylate lysine residues of proteins within a 10nm radius. Our approach involved the isolation of biotinylated proteins and the use of liquid chromatography mass spectrometry (LC MS/MS) to identify associated proteins in a quantitative manner (Figure 4a). Our experimental design compared expression of BirA*-DUX (full-length) in mESCs (which creates a 2C-like state) to expression of unfused BirA* alone (control) expressed in a 2C-like environment, in order to mimic a DUX-expressing 2C-like proteome in both conditions (Figure 4b). Here, we created clonal stable cell lines harboring our constructs, and expressed the proteins for 12hr with 0.25 μg/mL doxycycline and 5μM biotin. DESeq2 analysis of the sum intensity values from LC MS/MS revealed clustering between replicates (Supplementary Figure 4a) and a volcano plot revealed an enrichment in protein interactions for BirA*-DUX compared to BirA* alone, including those involved in chromatin de-repression such as the SWI/SNF (BAF) complex members SMARCC1 and SMARCA5 (Figure 4c, 4d). In addition to DESeq2 analysis, which is normally used for RNA-seq data, we applied the SAINTexpress package, which is designed for protein interactome analysis, to additionally analyze this dataset. DESeq2 appeared to be slightly less stringent than SAINTexpress analysis, with a larger number passing statistical threshold (p_adj_ <0.05) (Supplementary Figure 4b). SAINTexpress uses a statistical score called SAINTscore ranging from 0-1, with a significance threshold established as ≥ 0.73, which parallels an estimated protein-level Bayesian FDR of <0.05(34).

**Figure 4:**
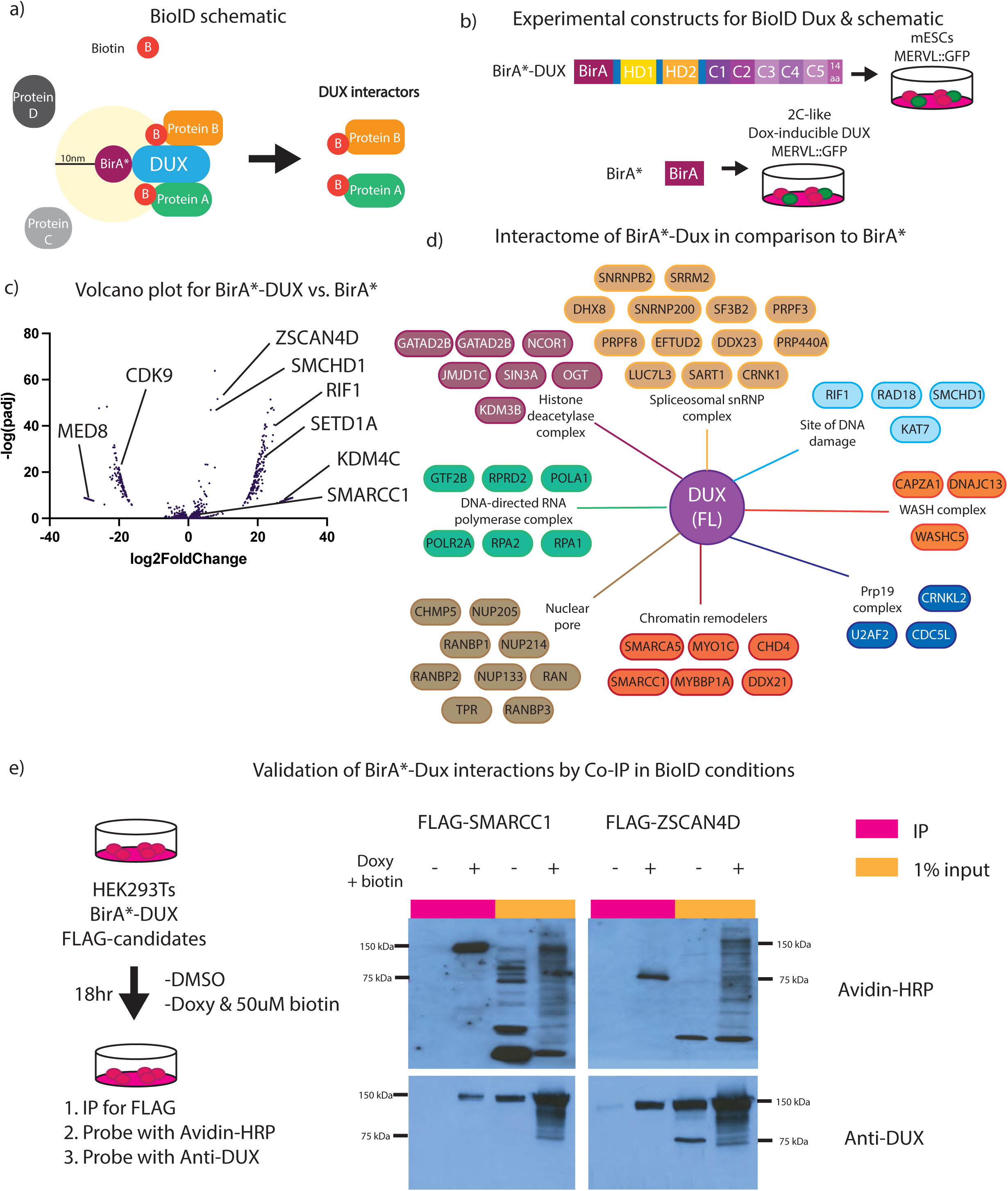
BioID for full-length DUX reveals interaction with proteins involved in chromatin de-repression. a. Experimental concept for BioID (proximity labeling assay) for BirA*-DUX to identify candidate DUX interacting proteins. b. Experimental setup for full-length DUX BioID. The control construct, BirA*-only, transfected into 2C-like mESCs, and BirA*-DUX, transfected into mESCs (n=2 biological replicates). c. Volcano plot using DeSeq2 analysis for BirA*-only vs. BirA*-DUX with log2FoldChange on the x-axis and -log(padj) on the y-axis (n=2 biological replicates). d. Interactome map of proteins enriched to interact with full-length DUX, vs. BirA*-only, split into specific cellular components and protein complexes (n=2 biological replicates). Full dataset available in Supplementary Table 1. e. (Left) Experimental setup for BioID-like experiment in HEK283T cells with BirA*DUX and flag-tagged candidates SMARCC1 and ZSCAN4D. (Right) Co-immunoprecipitation of FLAG and Western blot after 18-hour expression of transiently transfected FLAG-SMARCC1 or FLAG-ZSCAN4D and BirA*-DUX in HEK293Ts (n=3 biological replicates). Predicted protein size for BirA*-DUX is 125 kDa.

SWI/SNF subunit SMARCC1 and the transcription factor ZSCAN4D, two statistically significant hits from both analyses, were then selected for validation by testing for interaction *in vivo* via immunoprecipitation. We expressed N-terminally FLAG-tagged versions of SMARCC1 and ZSCAN4 in HEK293 cells co-expressing the BirA*-DUX fusion in the presence of 50 μM biotin (50μM biotin for BioID was empirically determined as the optimal dose in HEK293Ts). SMARCC1 and ZSCAN4 were immunoprecipitated with anti-FLAG conjugated beads, and analysis with an avidin-HRP blot showed they are both biotinylated and co-IP with DUX protein, validating BioID protein interactions (Figure 4e). Additionally, CHAF1A, an additional BioID hit, was also validated through FLAG-tag co-IP and DUX protein interaction, however KDM4C did not validate, suggesting that this hit was a false positive (Supplementary Figure 4c), and highlighting the importance of validating the BioID results.

Lastly, it has been previously reported that human DUX4 interacts with p300 (22,25), however p300 was not identified as a significant interactor in our dataset. This could be due to differences in the interactomes of human DUX4 and mouse DUX, or instead be technical, due perhaps to a lack of sensitivity to detect P300 peptides. To determine if mouse DUX similarly interacts with P300, we performed a BioID-like experiment in mESCs, which revealed p300 as modestly biotinylated, suggesting that a portion of p300 comes in close proximity with DUX, but was not annotated in the dataset, representing a possible false negative (Supplementary Figure 4d). In summary, our BioID experiment included chromatin-and transcription-related factors, a portion of which were validated using co-immunoprecipitation, which support prior observations of a strong impact of DUX occupancy on chromatin opening and transcriptional activity.

### Transcriptionally active DUX C-terminal repeat C3 binds to proteins involved in chromatin remodeling

Our work above presents the opportunity to identify proteins that interact differentially with a transcriptionally ‘active’ DUX derivative compared to a highly similar ‘inactive’ DUX derivative. Here, we separately N-terminally tagged our C1+14aa and C3+14aa derivatives with BirA*, to represent transcriptionally ’inactive’ and ‘active’ DUX constructs, respectively (Figure 5a). The experimental design and setup was identical to that described in Figure 4a, but the mESC line utilized bore a *Dux-/-* genotype. Furthermore, in place of the BirA* alone control, the inactive BirA*-C1 (BirA* fused to C1+14aa) was used as the control construct for comparison to the active BirA*-C3 (BirA* fused to C3+14aa).

**Figure 5:**
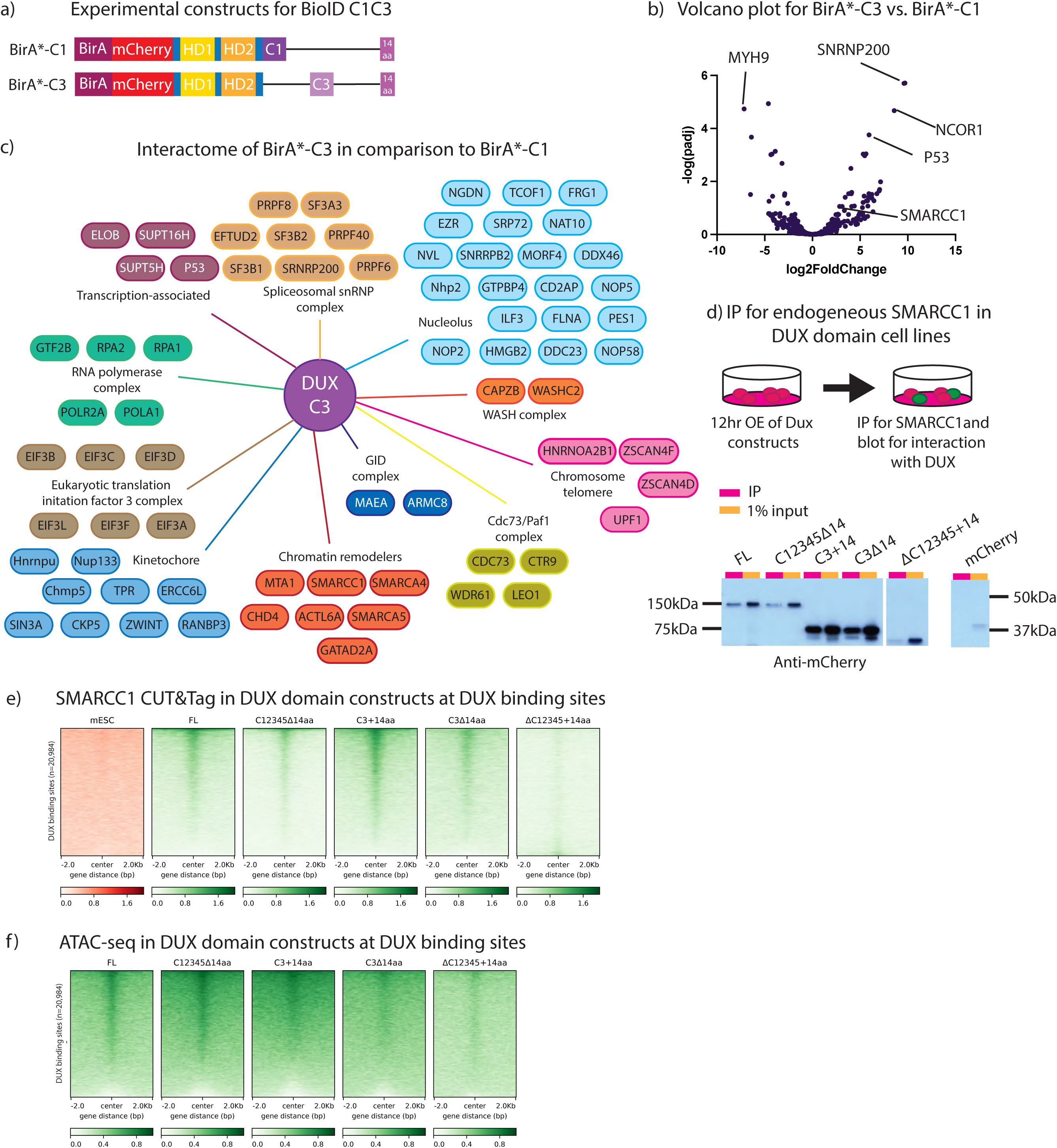
BioID for active C-terminal DUX repeat reveals interactors, including Smarcc1. a. Experimental constructs BioID of individual DUX repeats. Experimental constructs are N-terminally labelled BirA* linked to either mCherry-HDs-C1+14aa or mCherry-HDs-C3+14aa. b. Volcano plot using DeSeq2 analysis for BirA*mC-C1+14aa and BirA*mC-C3+14aa with log2FoldChange on the x-axis and -log(padj) on the y-axis (n=3 biological replicates). c. Interactome map of proteins enriched with BirA*mC-C3+14aa vs. BirA*mC-C1+14aa, split into specific cellular components and protein complexes (n=2 biological replicates). Full dataset available in Table 2. d. Co-immunoprecipitation of endogenous SMARCC1 after 12hr overexpression of DUX domain constructs in stable clonal cell lines illustration in Figure 2a (n=3 biological replicates). Predicted protein sizes are 100 kDa (FL), 95 kDa (C12345Δ14aa), 65 kDa (C3+14aa), 59 kDa (C3Δ14aa), 48 kDa (ΔC12345+14aa), and 25 kDa (mCherry). e. SMARCC1 CUT&TAG heatmap at DUX binding sites after 12-hour overexpression of Dux constructs illustrated in Figure 2a (n=2 biological replicates). f. ATAC-seq heatmap at DUX binding sites after 12-hour overexpression of Dux constructs illustrated in Figure 2a (n=2 biological replicates, except full-length DUX only has 1 biological replicate).

PCA and clustering analysis reveals that the three replicates of the ‘active’ C3 derivative group together and are more similar to each other (via clustering) than are the C1 replicates, and vice versa (Supplementary Figure 4e,f). Using SAINTexpress, we observed many more statistically-enriched (SAINTscore ≥ 0.73) protein interactions occurring with the DUX C3 ‘active’ repeat compared to the ’inactive’ C1 repeat. Specifically, we observed 12 proteins with the C1 derivative, and 154 proteins with C3 (Figure 5b). For C3 derivative interactors, GO-term analysis identified two overarching functional categories: chromatin and the nucleolus. Within the chromatin category, proteins within the NuRD, SWI/SNF and ISWI chromatin remodeling complexes were prominent (Figure 5c). Notably, proper formation of the nucleolus has been found to be required for Dux repeat locus silencing (35,36), and our data strongly suggests a direct physical interaction with nucleolar components, as DUX can bind to its own genetic locus (37).

To further investigate chromatin-related interactors of interest, we performed co-IPs for FLAG-tagged SMARCC1 (BAF155) (described above) using selected DUX domain derivatives. First, SMARCC1 binding to DUX appears only slightly diminished with the loss of the 14aa tail (Figure 5d). SMARCC1 also binds well to a DUX derivative containing both an active C3 repeat and the acidic tail (C3+14aa), and binding is largely maintained when the tail is removed (C3△14aa). Finally, a major reduction in SMARCC1 binding is observed when all repeats are removed and only the acidic tail remains. This suggests a hierarchy of interaction for DUX and SMARCC1: an ‘active’ repeat alone binds SMARCC1 whereas the acidic ‘tail’ alone does not bind SMARCC1, however the acidic tail slightly improves SMARCC1 interaction when active and inactive repeats are both present (full-length DUX).

Lastly, we wanted to determine whether the SWI/SNF (BAF) component SMARCC1 is recruited to DUX binding sites by DUX, and if so, which DUX domains confer this function. To address this, we performed CUT&Tag for endogenous SMARCC1 in the DUX derivatives and found, similar to our co-IP results, that SMARCC1 is recruited to DUX binding sites by DUX derivates that contain active C-terminal repeats but not by the 14aa tail (Figure 5e). Thus, binding and recruitment of SWI/SNF to DUX targets *in vivo* appears to rely on interaction with active DUX C-terminal repeats.

To continue investigating DUX interaction with SWI/SNF, we performed ATAC-seq in the DUX derivatives to determine whether SMARCC1 recruitment to DUX binding sites correlates with chromatin accessibility changes (Figure 5f, Supplementary Figure 4g). The results suggest a hierarchy of interaction similar to that observed in our SMARCC1 CUT&Tag experiments. At DUX binding sites, full-length DUX leads to increased opening, as do C12345△14aa and C3+14aa, the two derivatives which confer transcriptional activity. However, DUX derivatives which lack transcriptional activity or have decreased binding to SMARCC1 display a corresponding decrease in chromatin opening, though some opening does occur. This evidence further supports a combinatorial domain model for proper DUX protein interaction, occupancy, and target activation – revealing a major/required contribution from the C-terminal repeats and augmentation by the 14 aa C-terminal tail.

## Discussion

Embryonic genome activation (EGA) creates a transcriptome and proteome that replaces maternally-inherited components, reprograms chromatin and drives cleavage-stage processes (12). The transcription factor DUX is highly but transiently expressed during EGA (specifically the early 2-cell embryo stage) from a large locus containing ∼30 Dux repeats. DUX plays a key role in opening many promoters and enhancers, and in activating endogenous genes and repeat elements in the 2-cell embryo (37). DUX targets account for about 25% of the cleavage-stage transcriptome, and partly overlap with targets of OBOX4, underlying the observation that combined DUX and OBOX4 omission/depletion strongly affects embryo development (31). Although DUX targeting by its N-terminal homeodomains is well understood, how domains in the C-terminus of DUX contribute to chromatin reprogramming and transcription has remained largely unknown (23). Here, we conducted an analysis combined with proximity ligation to reveal the protein partners of DUX domains and the roles of DUX domains in chromatin remodeling and transcriptional activation.

First, we found that the mouse DUX C-terminal region contains five ∼100aa repeats (termed C1-5) followed by a 14aa acidic C-terminal tail, whereas DUX orthologs contain only one copy of this repeat, followed by a similar acidic tail. Our phylogenetic and DUX domain analyses suggest that two of the repeats (C3 and C5) perform a more active role than the ‘inactive’ C1/2/4 repeats. Interestingly, single ‘active’ mouse repeats show no transcriptional stimulation on their own, but show potent activity in combination with the acidic tail, essentially mimicking the domain-activity relationships of their human, rat or bovine DUX4 orthologs. Together, these observations suggest that additional DUX repeats in the mouse lineage may have derived from duplication of the original ‘active’ repeat, with subsequent sub-functionalization.

These observations raise two interesting questions – why does DUX have multiple repeats with different activities, and what distinguishes active from inactive repeats? Our work has localized the primary difference between active and inactive repeats to a small 6 aa region. Furthermore, active versus inactive repeats (in combination with the acidic tail) show very different profiles of protein interaction – with active repeats displaying more robust interaction with chromatin and transcription partners. Interestingly, we observe a cohort of genes (Figure 2f, cluster #3) that are silent with full-length DUX, silent with an ‘inactive’ repeat (C1), but active with a single ‘active’ repeat (C3). This raises the possibility that the ‘inactive’ repeats may serve to attenuate the activity of the active repeats under certain conditions or at certain locations. Of note, DUX binds to its own promoter to help activate a strong positive feedback loop; however, Dux activation is brief (half a cell cycle), with silencing accompanying the association of the large Dux locus with the nucleolar envelope(37,38). Interestingly, we observe high enrichment of nucleolar components with the ‘active’ C3 repeat, raising the possibility that the ‘active’ repeat may initially recruit powerful chromatin and transcription factors, and then subsequently contribute to localizing the Dux locus to a compartment that overcomes and/or reverses those associations to enable Dux locus silencing(35). Furthermore, interactions of DUX with histone deacetylase components were obtained, which remain to be fully explored in this silencing process. These observations provide new mechanistic information as well as opening new avenues for future exploration regarding the ability of DUX to function as a strong activator, while also helping to subsequently silence the repetitive Dux locus. Here, we envision that temporal and/or locus-specific post-translational modifications on DUX may mediate the temporal associations of DUX with target factors. Indeed, the presence of both ‘active’ and ‘inactive’ DUX repeats may help in coordination of these associations.

Regarding the functions of the DUX C-terminus, prior work of others has shown that the acidic C-terminal tail of DUX4 promotes transcriptional activation and interaction with the histone acetyltransferase p300(22). Our validation experiments confirmed those results with mouse DUX, and we extended our studies to explore the roles and relationships of the repeat regions, both alone and in combination with the acidic tail. Our examination of Dux domain constructs was extensive, and included RNA-seq, CUT&Tag, BioID, ATAC-seq and the ability to convert mouse ES cells to 2C-like cells. As DUX opens chromatin, and as three SWI/SNF complex components were present in our BioID results (SMARCC1, SMARCA4, ACL6A), these results prompted our direct assessment and validation of the recruitment of SWI/SNF complex (monitoring the core SMARCC1 subunit) to DUX binding sites (Figure 5c, Figure 6a).

**Figure 6:**
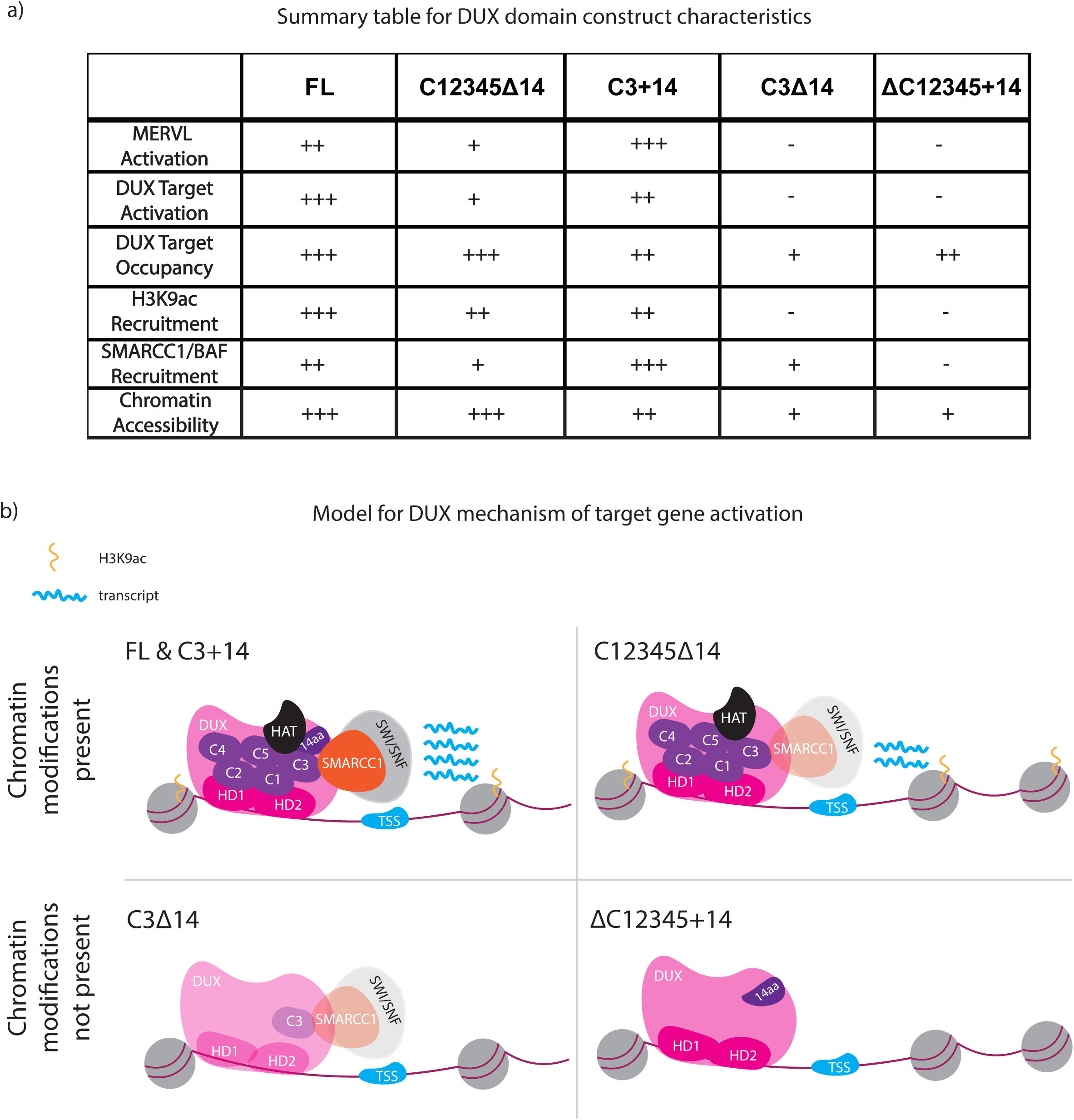
Model of DUX protein mechanism of activation based on C-terminal domain combinations. a. Summary table of DUX domain constructs (illustrated in Figure 2a) and their transcription, recruitment, chromatin modification, and protein interaction activities. b. Model for the contributions of DUX domains on DUX interactions and activity

An additional goal was to explore whether the functions and associations of the active repeats versus the acidic tail are largely separable/independent, or instead primarily additive or cooperative. First, regarding the activation of cleavage-specific genes (including MERVL), the target repertoire observe with full-length DUX is largely recapitulated with DUX derivatives containing the homeodomains and both active C-terminal repeats – even in the absence of the acidic tail. Thus, the tail accentuates activity, but is not strictly required for activity at the vast majority of targets. Regarding domain contributions to occupying DUX targets, all derivatives were able to bind DUX targets, though the absence of the acidic tail lowers occupancy if only one active repeat is present. This contrasts with the recruitment of H3K9ac activity; here, neither the acidic tail nor an active repeat is sufficient for recruitment; instead, either multiple repeats or repeat-tail combinations appear needed. For SMARCC1 (SWI/SNF), an active repeat is sufficient for recruitment, but recruitment is enhanced by the addition of the tail, whereas the tail alone appears ineffective. Finally, and of particular interest, a DUX derivative lacking repeat regions but containing the acidic tail (△C12345+14) occupies DUX targets, but does not activate targets – and does not recruit either H3K9ac or SWI/SNF complex. Therefore, the acidic tail appears to be effective only when combined with an active C-terminal repeat (Figure 6a). Overall, our data is consistent with largely cooperative functions for active repeat(s) and the acidic tail – as the absence of either compromises most functions and interactions – but places the C-terminal domains as more important domain in the hierarchy, as key factors like SWI/SNF complex are fully reliant on an active repeat for interaction. In keeping, our work predicts interactions with chromatin and transcription factors involving combined surfaces on repeats and the acidic tail (a two-domain model).

As previewed above, our proximity-labeling (BioID) revealed a variety of protein complex interactions, and we validated a subset of interactors with co-IP experiments, including particular chromatin and transcription-related factors. We note that certain interactions may be indirect and involve DUX association with component(s) involved in a larger multi-protein structures, such as the highly repetitive Dux locus (to which DUX binds(12,37)) in either its active or inactive form, and the association of the Dux locus with the nucleolus or other compartments. Thus, a complete understanding of the full repertoire of interactions will require future studies. However, beyond the factors noted above, we will emphasize two additional factors prominent in the BioID data that might be prioritized. First, the p53 interaction verifies prior work supporting p53 association with and activation of the Dux locus(37), and prompts examination of a direct interaction with DUX. Second, the presence in our dataset of SMCHD1, a cohesin-like protein known to help confer repression of the mouse Dux and human DUX4 repetitive loci, and to be mutated in a subset of FSHD patients, likewise prompts considerable future attention (39,40).

Taken together our work provides considerable new mechanistic information regarding the structure-function relationships and factor associations conducted by particular domains within the mouse DUX C-terminal region. Key findings include the characterization of ‘active’ repeat domains, their interaction with transcription and chromatin remodeling factors, and the cooperativity between DUX active repeats and the acidic tail in cofactor recruitment and transcriptional activation at DUX targets. In addition to factors for activation, factors related to repression and nucleolar association were also isolated, and future studies should help determine whether their temporal associations with *DUX* may help early embryos execute a quick ‘burst’ of Dux activation followed by repression at the Dux locus in 2-cell embryos.

## Specific author contribution

Experiments and analyses were conducted by C.M.S. with assistance from E.J.G. and S.C.S. C.M.S, E.J.G., S.C.S., and B.R.C. designed the study and C.M.S. and B.R.C. wrote the manuscript.

## Acknowledgements

We thank members of the Cairns laboratory and M.B. Chandrasekhran for inspiring discussions. We thank Brian Dalley for assistance with preparing RNA-seq libraries and for sequencing services, and T. Parnell for bioinformatic assistance. We give special thanks to Ross Tomaino at Taplin Mass Spectrometry Core (Harvard University) for mass spectrometry assistance for BioID experiments. Lastly, we thank Brian Patten at Novogene for Illumina sequencing services for CUT&Tag and ATAC-seq libraries.

## Materials Availability

Plasmids used in this study will be available by request.

## Data accessibility

RNA-sequencing, CUT&Tag, and ATAC-seq data are available through GEO GSE224300. Standard packages were used for RNA-seq, CUT&Tag, and ATAC-seq analyses (see “Materials and Methods”).

## Materials and Methods

### Amino acid alignment

DUX ortholog protein translations were aligned using Geneious Local Alignment (Smith-Waterman), Blosum62 cost matrix, gap open penalty of 12, and gap extension penalty of 4. Mouse embryonic stem cell culture Mycoplasma-free E14 mESCs were cultured on gelatin in 2i+LIF medium containing Gibco KO-DMEM with nonessential amino acids, 2-mercaptoethanol and dipeptide glutamine and were supplemented with 15% ESC-grade FBS, leukemia inhibitory factor (Thermo Fisher), 1 μM PD0325901 (Sigma-Aldrich) and 3 μM CHIR99021 (Sigma-Aldrich). Stable cell lines were blasticidin (Fisher Scientific, B12150-0.1) at 3 μg/mL.

### HEK293T cell culture

HEK293Ts were cultured in high-glucose growth media DMEM with nonessential amino acids, 2-mercaptoethanol and dipeptide glutamine and were supplemented with 10% FBS.

### Transfection of mESCs and HEK293Ts

Cells were transfected in Opti-MEM medium (Thermo Fisher Scientific, 31985070) using Lipofectamine 3000 (Thermo Fisher Scientific, L3000-015) for DNA transfections and the manufacturer’s recommendations. If transfection were to create stable clonal cell lines, a PiggyBac transposase was included in the transfection.

### Doxycycline treatment of cells

After cells have been plated for 24 hours (mESCs, or HEK293Ts), they were treated with doxycycline (0.25 ug/mL; Clonetech, 631311) for 12 h or 18 h to induce mCherry-DUX or DUX domain deletion construct expression, as indicated.

### Western blot

RIPA buffer (50 mM TRIS-HCl [pH 8.0], 150 mM NaCl, 1% NP-40, 0.5% sodium deoxycholate, 0.1% SDS) with 1X protease inhibitors was used to lyse mESCs. A 100X protease inhibitor cocktail consists of 0.03 mg/mL leupeptin, 0.14 mg/mL pepstatin A, 0.02 mg/mL chymostatin, 8.5 mg/mL phenylmethanesulfonylfluoride fluoride, 33mg/mL benzamidine and solubilized in ethanol. Lysed cell suspensions were then rotated for 30 min at 4°C and were further sonicated for 10 min with a 30-s on-off cycle in a Diagenode Bioruptor Pico device to solubilize chromatin-bound proteins. Protein lysates were quantified with the Bio-Rad (500-0001EDU) Bradford reagent and loaded on SDS-PAGE gel. After transfer to nitrocellulose membranes (VWR, 95040-108), membranes were blocked with TSBT (TBS with 0.1% Tween-20) and 5% milk and then incubated with antibodies overnight (mCherry ab167453, DUX (used in (37))and signals were detected using HRP-conjugated secondary antibodies and Western Lightning Plus-ECL, Enhanced Chemiluminescence (Perkin Elmer Health Sciences, NEL 105001EA).

### Flow cytometry

A Cytoflex LX instrument with lasers for 405 nm, 488 nm, 561 nm and 640 nm was used to quantify mCherry+ and MERVL–GFP+ mESCs. Samples were gated using forward FSA and side-scatter SSA to isolate cells from debris, and then double discrimination was performed using FSH Å∼ FSW and SSH Å∼ SSW.

### RNA-seq

1 million mESCs were grown for 24 hours on a 100mm dish and then induced with doxycycline (0.25 ug/mL) for 12 hours. Cells were washed two times with PBS, TRIzol (Thermo Fisher) was added to tissue culture wells and incubated for 1 min at room temperature. TRIzol/cells were added to Eppendorf tubes and RNA was extracted using the TRIzol extraction method. Libraries were generated according to the manufacturer’s instructions: polyA-selected RNA was isolated and libraries were prepared using the NEBNext kit (New England Biolabs, e7500s). Purified libraries were quantified on an Agilent Technologies 2200 TapeStation with a D1000 ScreenTape assay. The molarity of adaptor-modified molecules was defined by a qPCR with a KAPA Library Quantification kit (Kapa Biosystems). Individual libraries were normalized to 10nM, and equal volumes were pooled in preparation for Ilumina sequence analysis. Sequencing libraries (2pM) were chemically denatured and applied to an Illumina HiSeq paired-end flow cells with an Illumina cBot. Flow cells were then transferred to an Ilumina HiSeq 2000 instruct and sequenced in the 125-bp paired-end mode. RNA-seq reads were trimmed and filtered for quality using FastQC (version 0.5.0). Processed reads were aligned using STAR (version 2.7.2c). Differential expression analysis was performed using DESeq2 (version 3.11). Hierarchical clustering was performed using the R package ‘Complex Heatmap,’ with inputs being differentially expressed transcripts (DESeq FDR < 0.05) z-score normalized.

### GO term analysis

GO terms were identified using the Gene Ontology Resource (http://geneontology.org/).

### CUT&Tag

1 million mESCs were grown for 24 hours and then induced with doxycycline (0.25 ug/mL) for 12 hours. Cell were washed 1X with PBS and nuclei were isolated from mESCs using nuclear extraction buffer (20 mM HEPES-KOH [7.9], 10 mM KCl, 0.1% Triton X-100, 20% glycerol, 0.5 mM spermidine, 1X protease inhibitors (see above). ConA-streptavidin beads were paired with antibodies of interest 1:100 (mCherry ab167453, H3K9ac ab 32129, SMARCC1 ab172638) and 50,000 cells were added and incubated overnight in binding buffer (20 mM, HEPES [pH 7.9], 10 mM KCl, 1 mM CaCl_2_, 1 mM MnCl_2_). Secondary guinea pig anti-rabbit 1:100 (ABIN101961) was incubated for 1 hr at room temperature. pA-Tn5 (pre-loaded with Mosaic End A/B adaptors) was incubated for 1 hour at room temperature and used at 1:100. After washing steps and release from the beads, PCR was performed for 12 cycles using NEBnext High-Fidelity 2X PCR Master Mix (New England Biolabs, M0541L) (41). Samples were 1:1.3 AMPure XP (Beckman Coulter) purified and sent for sequencing. All libraries were sequenced on the Illumina HiSeq 6000 platform in 150-bp, paired-end format. Paired-end, raw read files were first processed by Trim Galore (Babraham Institute) to trim low-quality reads and remove adapters. Processed reads were then aligned to mm10 with the following parameters: (--end-to-end –very sensitive –no-mixed). CUT&Tag libraries were trimmed and filtered for quality using FastQC (version 0.5.0) and Picard (version 2.26.3). Processed reads were aligned using Bowtie2 (version 2.4.2) with the parameters: (--end-to-end –very sensitive –no-mixed) and indexed using Samtools (version 1.15). Duplicate samples were merged and normalized to control samples (mESCs for H3K9ac dataset and mCherry alone for mCherry and SMARCC1 datasets). DeepTools was used to generate heatmaps and profile plots using DUX binding sites as previously published (21).

### BioID

15 million BirA* mESC derivatives were plated on gelatin-coated 15-cm plates and recovered for 24-hours. 0.25 μg/mL doxycycline and 0.5 μM biotin (Sigma Aldrich, B4639) was added to cells and incubated for 18 hours. Cells were washed with ice cold PBS 2 times and 1 mL lysis buffer (2 mM Tris-HCl [pH 8.0], 137 mM NaCl, 1% NP-40, 2 mM EDTA, 1x protease inhibitors) was added to each plate and incubated for 1 min (33). Cells were scraped of off the plate and transferred to a 2-mL tube. Cells then rotated for 30 min at 4°C and were further sonicated for 10 min with a 30-s on-off cycle in a Diagenode Bioruptor Pico device to solubilize chromatin-bound proteins. Next, samples were centrifuged at 4°C at 12,000 rpm for 20 minutes and the supernatant was transferred to new tubes. Protein lysates were quantified with the Bio-Rad (500-0001EDU) Bradford reagent. For streptavidin pulldown, 2 μg of protein were added to pre-washed 50 μL streptavidin Sepharose bead conjugates (Cell Signaling Technology, 3419S) and rocked overnight at 4°C. Cells were pelleted and then washed 5X with wash buffer (10 mM TRIS [pH 7.4], 1 mM EDTA, 1 mM EGTA, 150 mM NaCl, 1% Triton X-100, 0.2 mM sodium orthovanadate, 1x protease inhibitors), boiled at 95°C in 30 μL 4X Laemmli buffer, and ran on a SDS-page gel until the samples were 1-inch into the gel. Protein bands/smears were cut out of the gel with razor blades and transferred to individual 1.7-mL Eppendorf tubes filled with ddH_2_0. Samples were overnight shipped on dry for liquid chromatography-tandem mass spectrometry sequencing (LC-MS/MS) to Taplin Mass Spectrometry Facility, Cell Biology Department, Harvard Medical School. Gel smears were reduced using 1 mM DTT in 50 mM ammonium bicarbonate for 30 min at 60°C and cooled, and 5 mM iodoacetamide in 50 mM ammonium bicarbonate was added for 15 min and incubated at room temperature. 5 mM DTT was added to quench the reaction, and then 5 ng/μL of trypsin (Promega) was added and samples incubated over night at 37 °C. Samples were then desalinated and resuspended in 10 μL of high-pressure liquids chromatography (HPLC) solvent A (2.5% acetonitrile, 0.1% formic acid). A reverse-phase HPLC capillary column was made by placing 2.6 μL M C_18_ spherical silica beads into a fused silica capillary (100-μL M inner diameter, ∼30-cm length). Next, each sample was loaded using a Famos autosampler (LC Packings) onto the HPLC capillary column. A linear gradient was created to elute peptides with increasing amounts of solvent B (97.5% acetonitrile, 0.1% formic acid). Eluted peptides were subjected to electrospray ionization couple to a mass spectrometer (Thermo Fisher Scientific) to produce tandem mass spectra whose b and y ion series patterns were compared to Sequest (Thermo Fisher Scientific) to establish protein identity. The sequencing data were filtered to 1 to 2% peptide false-discovery rate. To analyze BioID datasets, two approaches were taken. First, we utilized DESeq2 (version 3.11) to provide candidate genes for BirA*-DUX vs. BirA* and BirA*-C3 vs. BirA*-C1. Then, we used SAINTexpress (version 3.5.3) with parameter: (-L6). Formal candidates exhibited > 2 Log_2_FC and FDR < 0.05 and SAINT score > 0.74, for DESeq2 and SAINTexpress analyses respectively.

### Co-immunoprecipitation

1 million mESCs were grown for 24 hours on 100mm dishes and then induced with doxycycline (0.25 ug/mL) for 12 hours. Cells were trypsinized, rinsed once with PBS, and lysis buffer (2 mM Tris-HCl, pH=8.0, 137 mM NaCl, 1% NP-40, 2 mM EDTA, 1x protease inhibitors) was added to lyse cells. Cells then rotated for 30 min at 4°C and were further sonicated for 10 min with a 30-s on-off cycle in a Diagenode Bioruptor Pico device to solubilize chromatin-bound proteins. Protein lysates were quantified with the Bio-Rad (500-0001EDU) Bradford reagent. 3X FLAG Peptide-coupled beads (Sigma Aldrich, F4799-4MG) were pre-washed with lysis buffer, 1 mg of protein was added to 40 μL of pre-washed beads, and samples were incubated at room temperature for 4 hours. Beads were pelleted by low-speed centrifugation. Samples were washed 5X with wash buffer (10 mM TRIS [pH 7.4], 1 mM EDTA, 1 mM EGTA; pH=8.0, 150 mM NaCl, 1% Triton X-100, 0.2 mM sodium orthovanadate, 1x protease inhibitors [see above]), boiled at 95°C in 30 μL 4X Laemelli buffer, and run on an SDS-page gel for western blotting analysis.

### ATAC-seq

mESCs were grown for 24 hours and then induced with doxycycline (0.25 ug/mL) for 12 hours. Cells were washed 1X with PBS, pelleted, resuspended, and lysed in cold lysis buffer (10 mM Tris-HCl [pH 7.4], 10 mM NaCl, 3 mM MgCl2, and 0.1% IGEPAL CA-630). Nuclei were pelleted and resuspended in transposase buffer. The Tn5 (Illumina FC-121-1030), and the transposition reaction was carried out for 30 min at 37°C. After purification, the Nextera libraries were amplified for 15 cycles with NEBnext PCR master mix and purified with a Qiagen PCR cleanup kit. All libraries were sequenced on the Illumina HiSeq 6000 platform in 150-bp, paired-end format. Paired-end, raw read files were first processed by Trim Galore (Babraham Institute) to trim low-quality reads and remove adapters. Processed reads were then aligned to mm10 with the following parameters: (--end-to-end –very sensitive –no-mixed).

## Supplementary Figure Legends

**Supplementary Figure 1:**
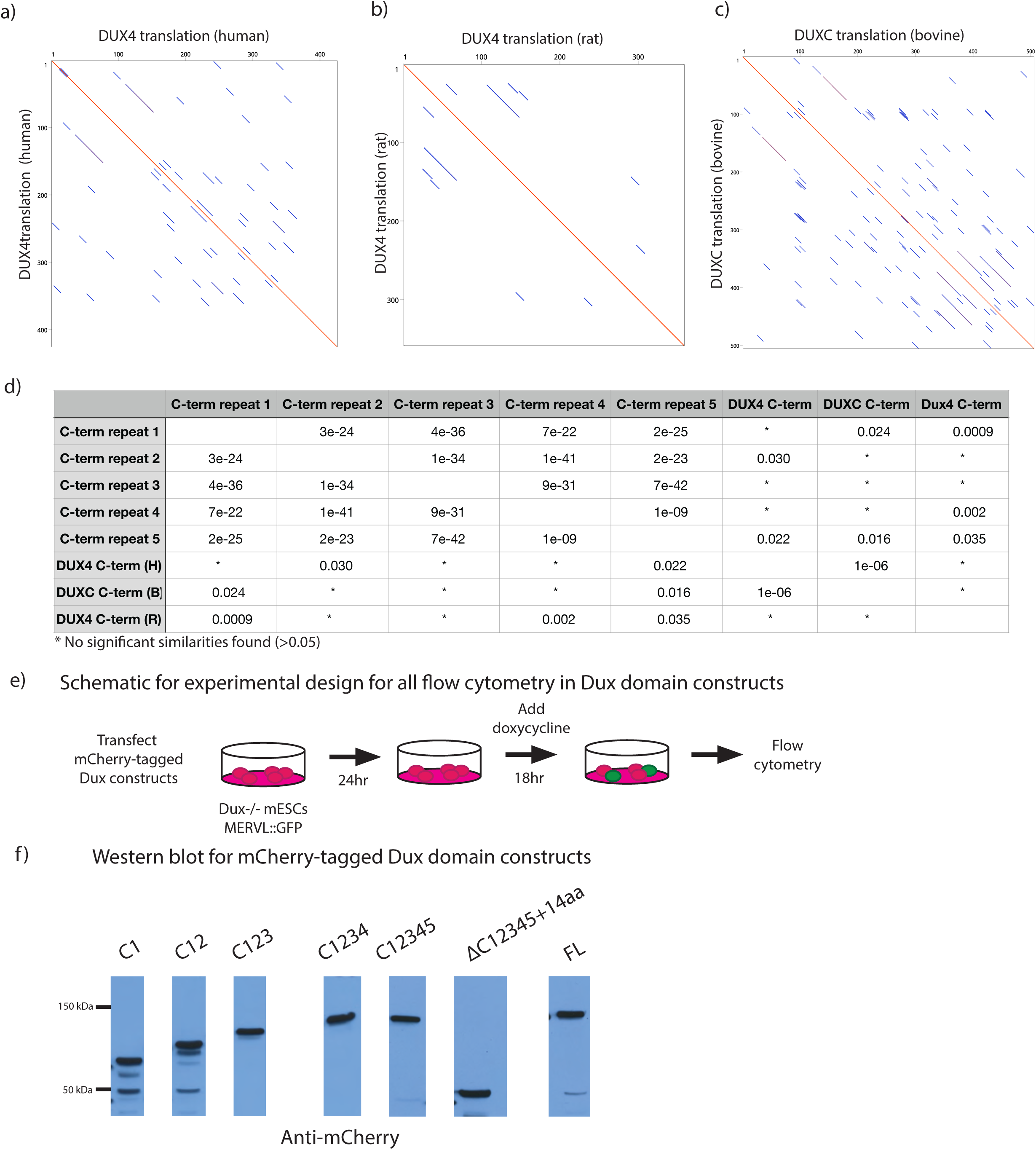
DUX orthologs do not contain a repeat structure similar to mouse DUX. a) Human DUX4 aligned to itself. b) Rat DUX4 aligned to itself. c) Bovine DUXC aligned to itself. d) Table with a pairwise comparison of e-values from a BLAST-based amino acid similarity comparing mouse DUX C-terminal repeats, the human DUX4 C-terminus, the rat DUX4 C-terminus, and the bovine DUXC C-terminus. Parameters are PAM250 matrix and gap costs of existence: 15 and extension: 3. *No significant similarities found (>0.05). e) Experimental design for expression of mCherry-tagged DUX domain constructs and flow cytometry of GFP% | mCherry tag expression. f) Western blot for mCherry tag after 18hr expression of C1, C12, C123, C1234, C12345. Predicted protein sizes are 59 kDa (C1), 69 kDa (C12), 79 kDa (C123), 89 kDa (C1234), 99 kDa (C12345), 48 kDa (ΔC12345+14aa), and 100 kDa (FL).

**Supplementary Figure 2:**
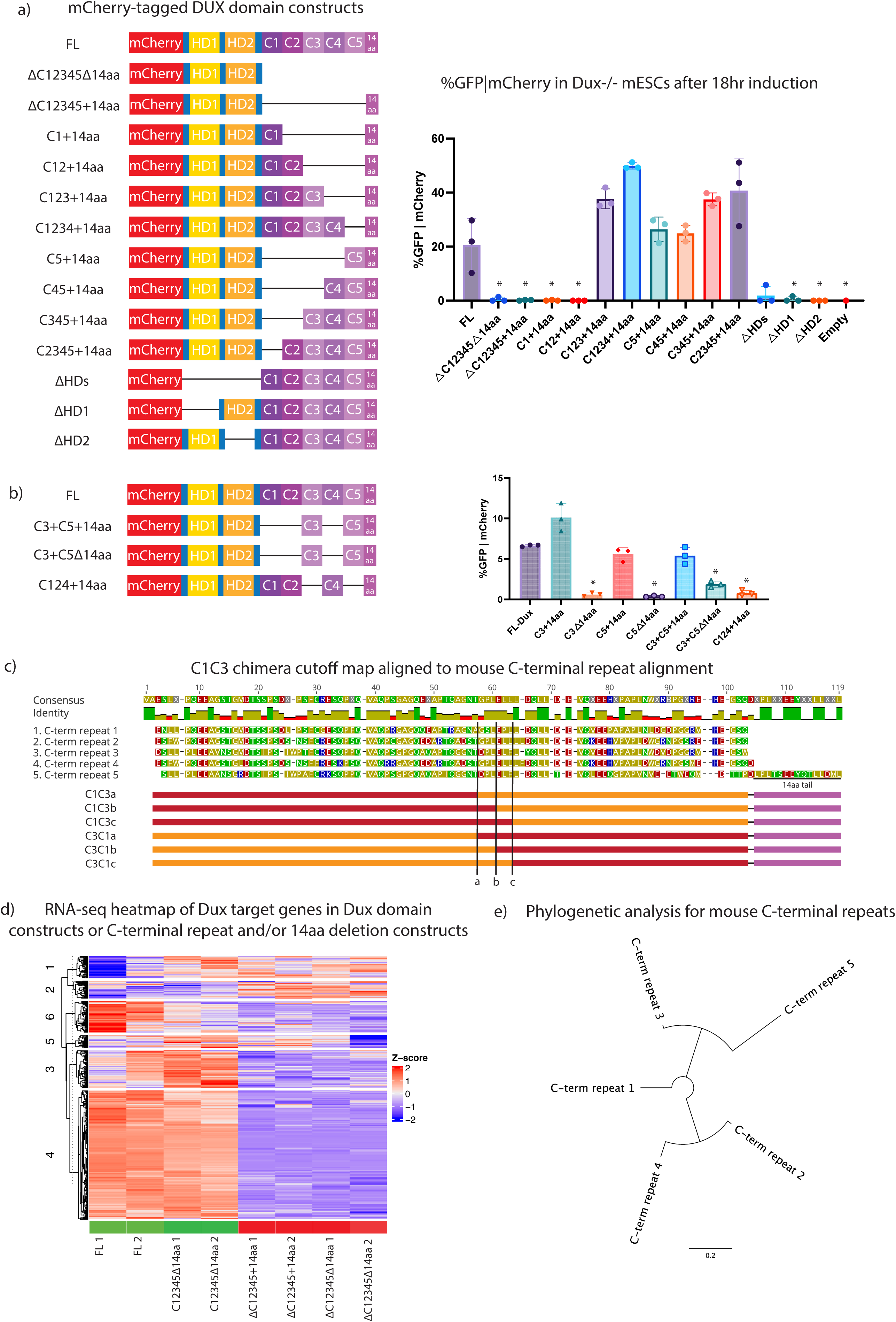
Supplementary Figure 2: DUX homeodomains are required for transcriptional activity. a) (Left) Schematic of constructs used for flow cytometry: mCherry-tagged full-length DUX (FL), ΔC12345+14aa, C12345Δ14aa, C1+14aa, C12+14aa, C123+14aa, C1234+14aa, C5+14aa, C45+14aa, C345+14aa, C2345+14aa, ΔHDs, ΔHD1, ΔHD2, and *Dux-/-* alone. (Right) Flow cytometry for MERVL::GFP reporter given mCherry expression in *Dux-/-* mESCs following 18hr expression of indicated constructs. *p-value < 0.05, student’s t-test. (n=3 biological replicates). b) (Left) Schematic of additional constructs used for flow cytometry: mCherry-tagged full-length DUX (FL), C3+C5+14aa, C3+C5Δ14aa, C124+14aa. (Right) Flow cytometry for MERVL::GFP reporter given mCherry expression in *Dux-/-* mESCs following 18hr expression of indicated constructs. *p-value < 0.05, student’s t-test. (n=3 biological replicates) c) Schematic of C1C3 chimera DUX constructs for amino acid cut offs for C1C3a, C3C3b, C1C3c, C3C1a, C3C1b, and C3C1c aligned to the 5 C-terminal repeats d) Heat map of RNA-seq at DUX target genes (n=456) (21) from 18hr expression of DUX domain constructs of FL, C12345Δ14aa, C12345+14aa, C12345Δ14aa(n=2). Colored rectangles above samples denote inability to activate MERVL::GFP reporter (red) and ability to activate MERVL::GFP reporter (green). e) Phylogenetic analysis of the five mouse DUX C-terminal repeats. Line length unit is substitutions per base.

**Supplementary Figure 3:**
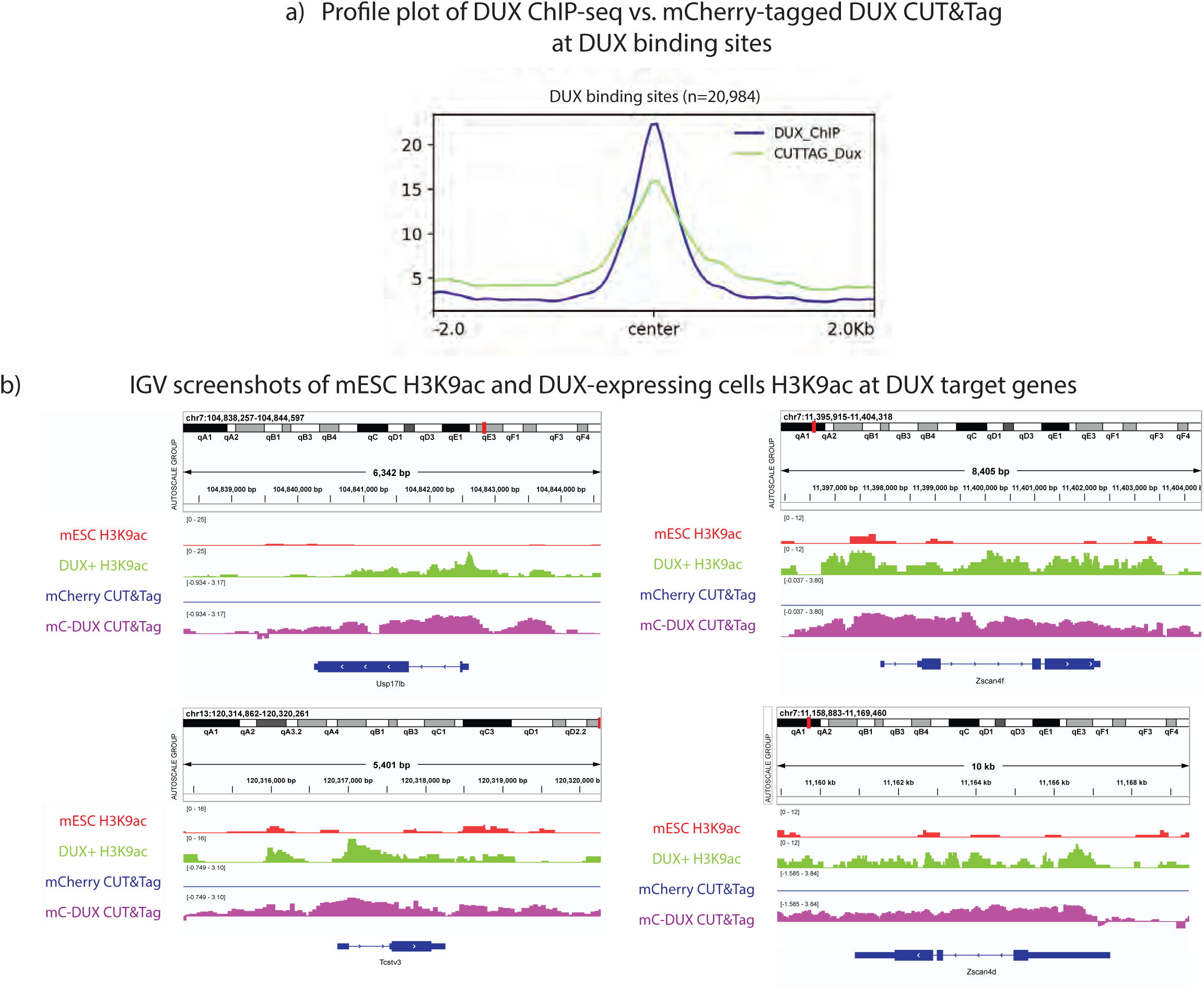
DUX CUT&Tag data strongly resembles DUX ChIP-seq, and DUX binding sites gain H3K9ac upon DUX expression. a. Profile plot of mCherry-FL DUX CUT&TAG centered at DUX binding sites after 12 hour expression vs. HA-DUX ChIP-seq (n=2 biological replicates) (12) b. IGV screenshots at TCSTV3, ZSCAN4F, USP17LB, and ZSCAN4C (DUX targets genes) for H3K9ac CUT&Tag in mESCs and mCherry-DUX 12hr expression and mCherry CUT&TAG for mCherry tag alone and mCherry-DUX (n=2 biological replicates)

**Supplementary Figure 4:**
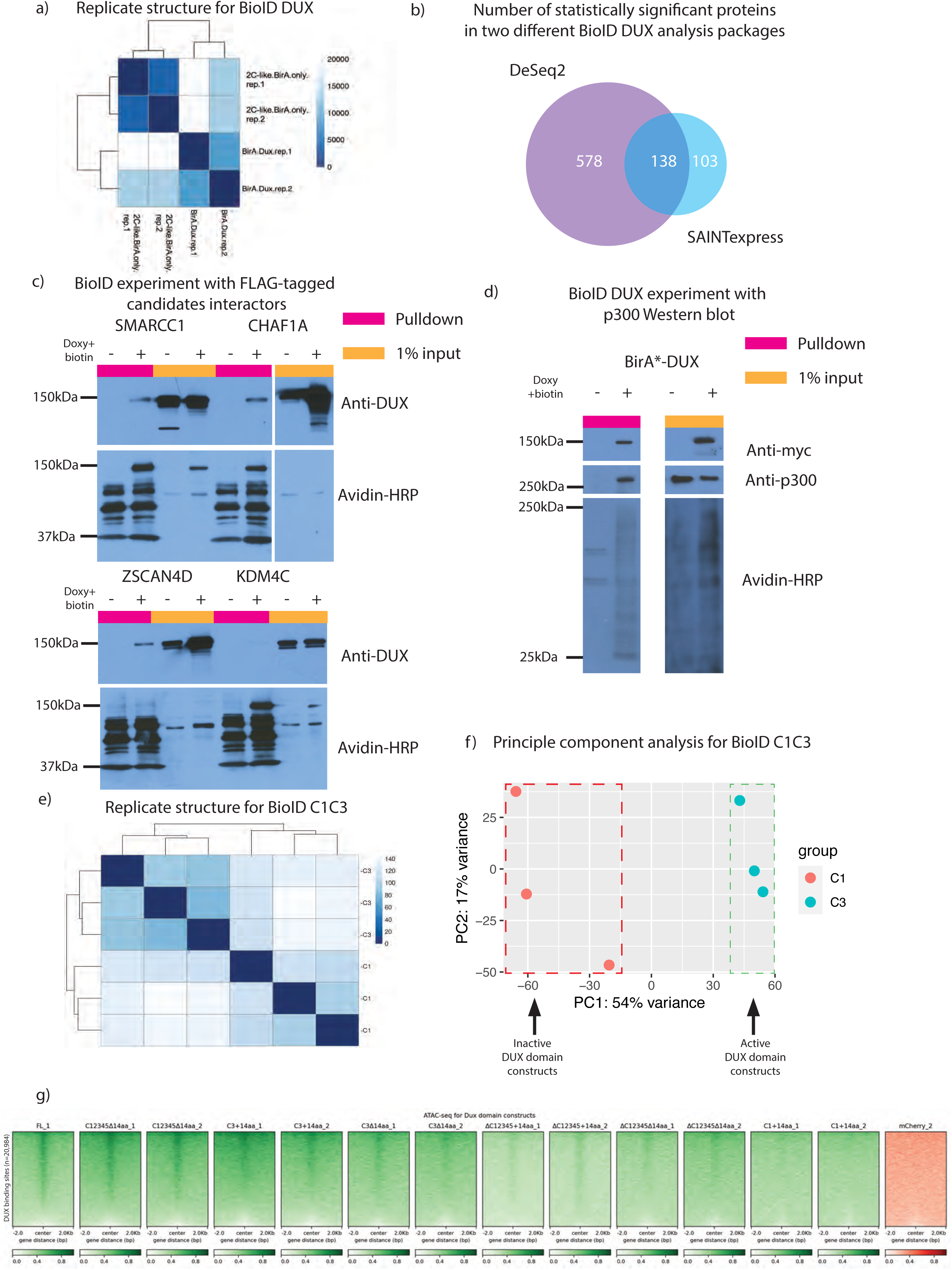
The proximity-labeling assay BioID identifies DUX protein interactors. a) BioID DeSeq2 analysis replicate structure of BirA*-only and BirA*-DUX duplicates for mass spectrometry datasets. b) Venn diagram of statistically significant enriched proteins for BirA*-DUX interaction for DeSeq2 and SAINTexpress c) Streptavidin pulldown and Western blot after 18-hour BioID induction in clonal stable BirA*-DUX mESC line expressing FLAG-tagged SMARCC1, CHAF1A, ZSCAN4D, or KDM4C with blots for DUX and FLAG-tag (n=3 biological replicates). KDM4C was a BioID hit, but does not interact with DUX, indicating a possible false positive. Predicted protein size is 125 kDa (BirA*-DUX). d) Streptavidin pulldown and Western blot after 18-hour BioID induction in clonal stable BirA*-DUX mESC line, with blot for N-terminal myc-BirA*-DUX, P300, and Avidin-HRP (n=2 biological replicates). Predicted protein sizes are 125 kDa (myc-BirA*-DUX) and 263 kDa (p300). e) BioID individual repeat DeSeq2 analysis replicate structure of BirA*-C1 and BirA*-C3 triplicates for mass spectrometry (n=3 biological replicates) f) Principle component analysis of DeSeq2 analysis for BirA*mC-C1+14aa and BirA*mC-C3+14aa with PC1: 54% variance and PC2: 17% variance g) ATAC-seq heatmap at DUX binding sites after 12-hour overexpression of duplicate Dux constructs illustrated in Figure 2a and C1+14aa

